# Mapping the Molecular Landscape of Thyroid Neoplasms: A Comprehensive Proteomic and Phosphoproteomic Analysis Across Tumors of Follicular Origin

**DOI:** 10.1101/2024.11.06.622203

**Authors:** Jonas Bossart, Kiarash Tajbakhsh, Olga Stanowska, Achim Weber, Robert Zboray, Aurel Perren, Marija Buljan

**Author notes:** Corresponding authors: Marija Buljan, Aurel Perren, Robert Zboray.

## Abstract

Thyroid nodules are a widespread phenomenon, with follicular cell-derived thyroid tumors being the most prevalent type of endocrine tumor, spanning from benign through low grade malignant to aggressive neoplasms with dismal prognosis. In clinical practice, histopathological criteria are primarily used to determine malignancy and aggressiveness. Therefore, accurate classification may result in surgical procedures for diagnostic reasons, associated with an imbalanced risk/benefit ratio. In recent years, the use of integrated proteomic approaches has proven valuable in expanding the molecular understanding of thyroid neoplasms, with implications in classification, yet remains understudied in divergent thyroid nodules. Here we show the delineation of subtype-specific and malignancy-dependent molecular characteristics through integrative proteomic and phosphoproteomic analysis of 53 human thyroid tissues, encompassing five frequent benign and malignant tumors. We found that the (phospho)-proteomic profiles enable a clear stratification of malignant and benign thyroid tissues. The method also performs well in delineating follicular adenoma (FA) and follicular thyroid carcinoma (FTC) samples. Beside the dysregulation of cell cycle control, apoptosis, and metabolic reprogramming associated with tumor development and malignancy, we further report increased alterations within the well-established oncogenic RAS/BRAF/MAPK and AKT/MTOR signaling pathways, which, contrary to the prevailing paradigm, did not clearly differentiate between FTC and papillary thyroid carcinoma (PTC). In addition, activities of ATM, PLK2-3, and GRK5-6 kinases were predicted to be strongly upregulated in malignant subtypes. Together, this study provides an in-depth insight into molecular changes in different thyroid tumor subtypes. These findings highlight the potential of integrated proteomic approaches to refine our understanding of complex diseases like cancer. As such, they offer a pathway to more precise diagnostic and personalized treatment strategies.

## Introduction

With a prevalence of up to 68%, thyroid nodules are a widespread endocrine phenomenon^1^, whereby the vast majority are identified as benign lesions^2^. However, the reliable stratification of follicular cell-derived thyroid tumors, particularly in the absence of clear hyperplastic changes or characteristic nuclear features of papillary thyroid carcinoma (PTC), possess a challenge^3^ that is still not unanimously resolved. The World Health Organization (WHO) has defined a classification system that categorizes follicular-derived thyroid tumors according to prognostic risk categories into benign, low-risk, or malignant neoplasms^4^. The benign tumors include, among others, oncocytic adenoma (OA) and follicular adenoma (FA), with OA consisting of more than 75% oncocytes, i.e. large cells with granular cytoplasm due to accumulation of dysfunctional mitochondria^4,5^. The malignant neoplasms are frequently well-differentiated. However, the rare but high-grade poorly differentiated thyroid carcinoma (PDTC) subtype is associated with aggressive features^4^ and can arise *de novo* or evolve from well-differentiated thyroid carcinoma^6^. Among well-differentiated thyroid carcinomas, PTC and follicular thyroid carcinoma (FTC) represent the most frequent histological subtypes^7^. Despite the shared follicular origin, PTC and FTC differ in terms of their nuclear features, with papillary-like changes being only found in the former. Treatment options available for progressed thyroid neoplasms include surgery, radioactive iodine (RAI) ablation, thyroid stimulating hormone (TSH) suppression, and immunotherapies^8–10^.

The prognostication and choice of cancer treatment are largely guided by clinical and histopathological features^11^. Molecular analyses have confirmed the important differences between FTC and PTC. The mutually exclusive RAS Q61R (FA: 10%, FTC: 22%, PTC: 8%) and BRAF V600E (FTC: absent, PTC: 24-51%) mutations are frequently observed in FTC and PTC, respectively^12–14^. Such molecular insights have proven valuable for refining and adjusting the guidelines for the diagnosis of different thyroid tumors in the past^4^. An integrated genomic characterization conducted by the Cancer Genome Atlas (TCGA) research network described substantial differences in BRAF V600E– and RAS-like PTC on transcriptome level^14^. Accordingly, BRAF V600E PTC preferentially signal via MAPK, while RAS-like PTC signal both via MAPK (albeit to a lower degree^15^) and PI3K^14^. In line with the advanced understanding the modern WHO classification system also considers differences in driver mutations. As such, the encapsulated invasive follicular PTC variant, which in contrast to classical PTC harbors a RAS and not classical BRAF V600E driver mutation, is molecularly closer to FTC^4^, and such carcinomas are coined as RAS-like cancer.

There are other common alterations affecting the PI3K/AKT/MTOR signaling axis, which is mutated much more frequently in PDTC (11%) and anaplastic thyroid carcinoma (ATC: 39%) than in PTC (1.4%)^16^ There is clinical evidence that activation of this signaling axis promotes proliferation and dedifferentiation of thyroid tumors of follicular origin through the inhibition of anti-tumorigenic mechanisms like apoptosis^17^. Accordingly, co-occurring mutations within the PI3K/AKT/MTOR pathway in BRAF V600E mutant PTC cases have a negative impact on patients disease-specific mortality^18^. In PDTC, BRAF V600E and RAS mutations occur with the same frequency, however, the mutational status represents the initial tumor they evolved from (FTC-derived PDTC are mostly RAS-like and PTC-derived PDTC BRAF V600E-like)^19^. Frequent mutations further include fusions of RET and NTRK1 with different partner genes. While RET rearrangements are predominantly observed in PTC (20%), those involving NTRK1 are not particularly associated with specific histological features, i.e. thyroid tumor subtypes^20^. Additionally, TERT promoter mutations (benign: 0%, FTC: 17%, PTC: 11%, PDTC: 43%)^21^ and nuclear deregulation of the proto-oncogene MYC (FTC: 25%, PTC: 24%, ATC: 76%)^22^ have been associated with both tumor aggressiveness and dedifferentiation.

Besides the investigation of specific cancer drivers, global transcriptomic and proteomic studies have also enabled the characterization of other novel molecular characteristics within key biological pathways. Among others, an integrated transcriptional analysis of patient samples has connected CREB3L1 with the progression of PTC to ATC^23^. Further, a transcriptome analysis of 671 PTC patients in the TCGA atlas has shown that high CREB3L1 expression negatively predicts patients’ overall survival^23^. Moreover, an enhanced lipid metabolism has been observed in PTC through a multi-omics and proteomics analysis, respectively^24,25^. The relevance of this observation was further underlined by the significant association of the transcript levels of three key enzymes within the lipid metabolic pathway (LPL, FATP2, CPT1A) with the tumor, node, metastasis (TNM) staging^24^. Recently, researchers demonstrated the capability of a 19-protein panel derived from over a thousand thyroid nodule proteomes to differentiate between benign (normal tissue/lymphocytic thyroiditis, multinodular goiter, FA) and malignant (FTC, PTC) entities^26^. In contrast to classical proteomics, which aims to comprehensively characterize all proteins within a sample, phosphoproteomics specifically focuses on the analysis of protein phosphorylation states. Recent technological advances, including the improved sensitivity of employed mass spectrometry (MS) techniques and phosphopeptide enrichment protocols, have facilitated the application of phosphoproteomics to clinical samples. Phosphoproteomics analysis has recently been applied to PTC^27^, where it added an additional layer for the characterization of stratified groups according to molecular subtypes and recurrence risks. This emerging technology holds promise in supporting the discovery of novel disease drivers and treatment targets, as it has the potential to more precisely illuminate aberrant cellular signaling networks.

In this study, we determined and compared the global protein expression and phosphorylation levels of different thyroid neoplasms of follicular origin. We applied an untargeted liquid chromatography (LC)-tandem MS approach to a multi-centric biobank cohort containing a total of 53 thyroid tissues, including 8 FA, 8 OA, 7 FTC (with two additional biological replicate samples from the same resections, i.e. 9 FTC samples were measured in total), 16 PTC, 8 PDTC, and 6 healthy control tissues. This approach enabled the identification of profound subtype-specific and malignancy-dependent molecular alterations affecting key biological and disease pathways and allowed us to connect changes in the levels of phosphorylated residues with altered kinase activities. As such, we were able to align changes in the (phospho-)proteome with existing knowledge related to oncogenic signaling axes and to expand the molecular characterization. Our integrative analysis benefits from improved stratification between thyroid tumor subtypes, particularly between FA and FTC. Interestingly, one FA sample clustered with the FTC group, and re-evaluation revealed borderline angioinvasion, suggesting that the tissue may be precancerous and not yet fully invasive. The analysis further highlighted the importance of the RAS/BRAF/MAPK and AKT/MTOR signaling pathways that were found more strongly dysregulated in malignant (FTC, PTC, PDTC) thyroid tumors. However, we observed that PTC, which is commonly considered a BRAF-like tumor, was also associated with increased NRAS protein expression, while FTC, which is considered RAS-like, also showed enhanced signaling via BRAF. Additionally, we predicted increased activities of the ATM, PLK2-3, and GRK5-6 kinases in different malignant thyroid tumors, identifying them as potentially interesting therapeutic targets. The molecular differences identified in this study, encompassing protein expression levels, phosphorylation patterns, and inferred kinase activities, provide a rich resource and have the potential to serve as candidates for the development of a more robust tumor classification system as well as for the formulation of improved personalized treatment regimens.

## Results

### Malignant thyroid neoplasms exhibit distinct and subtype-specific global proteome and phosphoproteome profiles that distinguish them from healthy thyroid tissue

This study encompasses the global MS-based proteome and phosphoproteome profiling of six thyroid tissue subtypes collected at two Swiss university hospitals (Zürich and Bern), measured in three individual batches, for which additionally BRAF V600E and RAS Q61R mutations were screened by immunohistochemistry (IHC) (**Fig. 1A**, **Table S1**). The 55 thyroid samples derived from 53 thyroidectomies were measured with a label-free quantification (LFQ) protocol in data-dependent acquisition (DDA) mode and the measured peptides were matched to the human proteome using the MaxQuant software tool^28,29^. The subsequent data processing resulted in the quantification of a total of 2’540 proteins and 5’618 phosphopeptides mapped to 1’850 phosphoproteins. We imputed missing data stepwise following concepts of the PhosR^30^ method, which considers the fraction and values of quantified proteins or phosphopeptides in each analyzed condition (see Methods). The basic rationale is that if most samples in a condition (i.e. more than half) have a specific entry measured, the missing values in the corresponding condition will be imputed close to the mean value of the individual intensity distribution of the samples, but if the majority is not measured, the missing values will be imputed from the lower end of the distribution. This strategy is expected to strengthen the clustering of biological groups compared to generic imputation methods (i.e. k-nearest neighbor or background imputation) and to lead to a higher number of significant hits in the downstream differential expression and pathway analyses^30^. Following imputation, the data was median-centered and batch corrected. Next, we applied hierarchical clustering (HC) (**Fig. 1B-C**, **Fig. S1A-D**) and principal component analysis (PCA) (**Fig. 1D-E**) to the processed data and observed a clear stratification of malignant and healthy control thyroid tissues. As expected based on previous proteomic studies^26,31^, the subtypes within the benign (Healthy, FA, OA) and malignant (FTC, PTC, PDTC) groups could not be separated completely. However, the applied protocol specifically enabled us to distinguish FA and FTC, offering the unique opportunity to assess differences between the two follicular tumors with different prognosis, as discussed in a separate chapter further below. This is of particular interest as the determination of follicular pattern thyroid tumors still remains one of the main diagnostic challenges^26^. Furthermore, the clustering of two biological replicates, which were measured in different batches, indicated that batch effects did not predominate the signals. Quantified phosphorylation sites included 4924 (87.6%) phosphoserine, 630 (11.2%) phosphothreonine, and 64 (1.1%) phosphotyrosine residues (**Fig. 1F**), following an expected pattern for the applied enrichment strategy^32^. Intra-class Pearson correlation analysis of global proteomes showed highest similarity level within healthy tissues (**Fig. 1G**). Pearson correlation coefficients were slightly decreased in benign FA as well as well-differentiated FTC and PTC samples, however, a much lower correlation coefficient was observed within PDTC and OA samples. For PDTC, this can be explained by the dedifferentiated cell states, high tumor heterogeneity, and more aggressive tumor behavior^33,34^. The low correlation among OA samples might appear counterintuitive. However, the high prevalence of mutations in the mitochondrial genome, which is known to cause compensatory mechanisms inducing the oncocytic phenotype^35^, could explain the high degree of heterogeneity.

**Figure 1:**
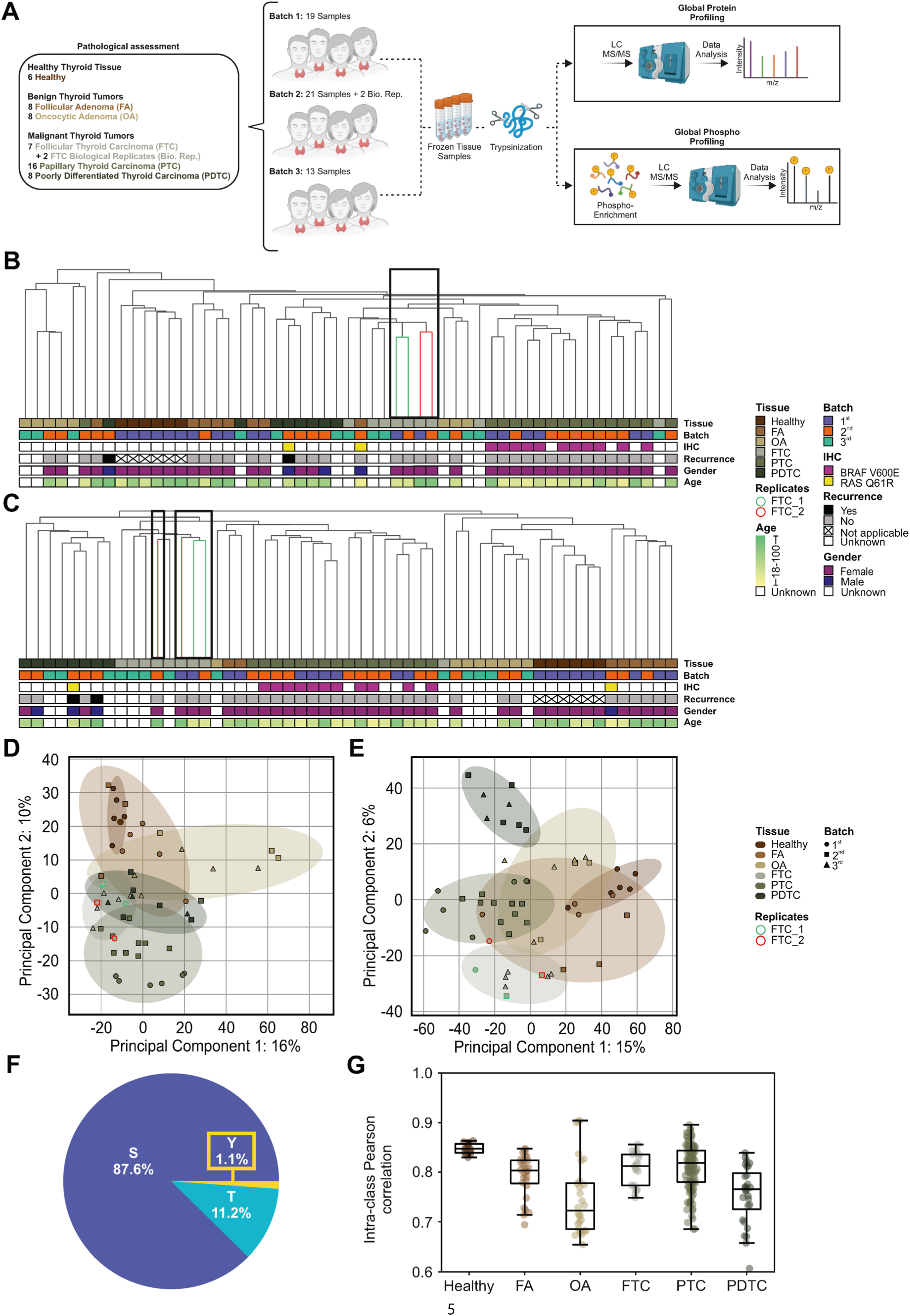
Proteomic and phosphoproteomic analysis of different thyroid neoplasms of follicular origin. (**A**) The experimental approach includes the measurement of 55 samples derived from 53 thyroid tissues. **B, C** Hierarchical clustering (HC) dendogram with information on tumor subtype, batch, immunohistochemistry (IHC) staining, recurrence, gender, and age of (**B**) 2’540 proteins and (**C**) 5’618 phosphopeptides. Intensities of individual entries are not displayed (see **Fig. S1B-C**). Biological replicates of two FTC samples (FTC_1 marked green and FTC_2 red in the dendogram) are highlighted with black boxes. **D, E** Principal component analysis (PCA) clustering including 95% confidence ellipses based on the global (**D**) proteome and (**E**) phosphoproteome. (**F**) Distribution of measured phosphorylation sites. (**G**) Boxplot of Intra-class Pearson correlation coefficients of proteomic data by thyroid tissue type.

### Differential expression and phosphorylation status of several cancer drivers show subtype-specific association with thyroid tumors

To obtain insights into the quantity and identity of differentially regulated molecular entities among the thyroid neoplasms, we conducted differential expression analyses (DEA) based on the quantitative values of a total of 2’540 proteins and 5’618 phosphopeptides. We compared protein levels in tumor tissues with those in healthy control and identified a total of 182, 406, 230, 599, and 360 differentially expressed entries in FA, OA, FTC, PTC, and PDTC tissues, respectively (moderated *t*-test, false discovery rate (FDR) < 0.05 and absolute Log2 FoldChange (FC) ≥ 1, **Fig. 2A**, **Table S2**). The observed increase in the number of differentially expressed proteins from FA (182) to FTC (230) and PDTC (360) is in line with the proposed mechanism of accumulated genetic alterations during thyroid tumor progression^36^. The larger sample size of PTC samples (N = 16) compared to other analyzed subtypes and the associated increase in statistical power contributes to the highest number of identified differentially expressed proteins (599) in this subtype, which would intuitively be expected in PDTC due to the advanced dedifferentiation. Furthermore, the previously mentioned prevalence of molecular alterations within the mitochondria in OA cells^35^ partially explains the high number of differentially expressed proteins in this subtype (406), as 172 out of the 406 significant proteins (42%) are associated with the Gene Ontology Cellular Compartment (GOCC) term *mitochondrion* (GO:0005739). The same GOCC term is far less prevalent in differentially expressed proteins in other thyroid neoplasms (FA: 24%, FTC: 19%, PTC: 21%, PDTC: 19%). By applying the same statistics and threshold values on the level of phosphopeptides, we found that the total number of significantly differentially phosphorylated peptides was markedly lower in benign (FA: 589, OA: 595) than in malignant (FTC: 1’287, PTC: 2’212, PDTC: 1’940) thyroid tumors (**Fig. 2B**, **Table S3**). The numbers of significant (phospho-)proteome alterations across the analyzed tumor subtypes are illustrated in Volcano plots (**Fig. 2C-D**, **Fig. S2**). Overall, this indicates that the more aggressive forms of thyroid tumors are often accompanied with the increased number of alterations in the levels of the expressed proteins and phosphopeptides. An exception to this pattern is OA, which shows an increased number of differentially expressed (but not phosphorylated) proteins presumably due to increased compensatory mitochondrial biogenesis and production of mitochondrial proteins.

**Figure 2:**
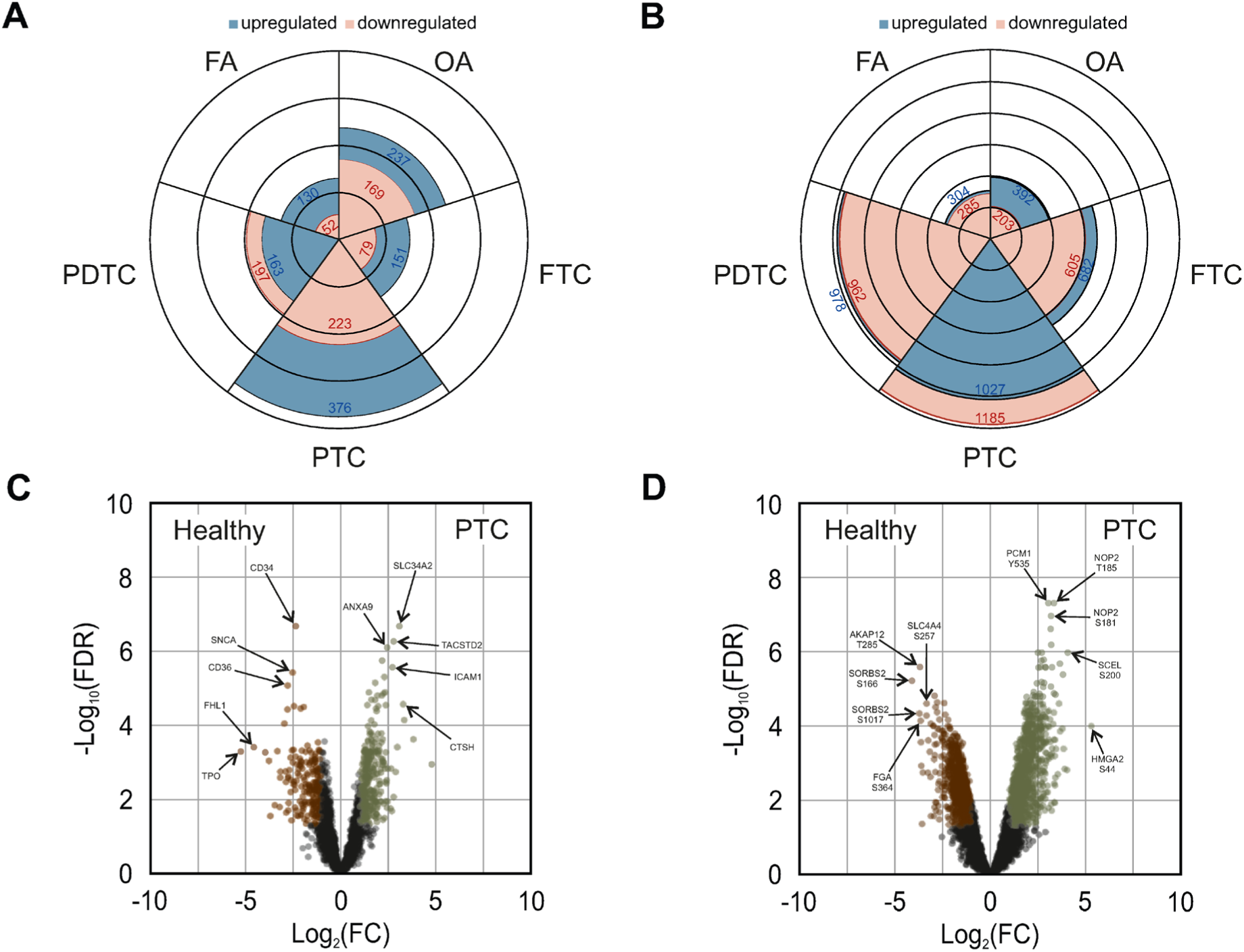
Thyroid tumor subtype-specific differential expression and phosphorylation of proteins. **A, B** Rose charts showing the numbers of upregulated (blue) and downregulated (pink) (**A**) proteins and (**B**) phosphopeptides of thyroid tumors compared to healthy thyroid controls. Applied thresholds include FDR < 0.05 and absolute Log_2_(FC) ≥ 1. **C, D** Volcano plots of quantified (**C**) proteins and (**D**) phosphopeptides comparing PTC with healthy thyroid controls. Colored entries represent significant hits with FDR < 0.05 and absolute Log_2_(FC) ≥ 1. Top five up– and downregulated hits were annotated based on their ranking according to –Log_10_FDR × Log_2_(FC).

We further evaluated if the differentially regulated proteins had established roles in thyroid cancer or were pointing to a dysregulation of kinases, transcription factors (TFs), or epigenetic factors (EFs), i.e. major regulators of cellular processes. For this, we assessed if any of the differentially expressed entries were annotated with respective roles or functions in the thyroid cancer pathway (hsa05216) available through the Kyoto Encyclopedia of Genes and Genomes (KEGG)^37^ and the databases from PhosphoSitePlus^38^, TRRUST^39^, as well as EpiFactors^40^. In addition, we assessed alterations within specific phosphorylation sites (that are not simply explained by altered protein expression levels, see Methods) with effects on protein stability and function according to UniProt^41^ annotations. In total, we observed significant alterations in the expression or phosphorylation levels of 4 proteins and 8 phosphoproteins with a known role in the thyroid cancer pathway, 17 kinase proteins and 64 kinase phosphoproteins, 18 TFs and 116 phosphorylated TFs, and 43 protein EFs and 192 phosphorylated EFs (**Table S2-S3**). Activating mutations in the RAS, BRAF, and RET genes, as well as within the main regulators of the MAPK signaling pathway are considered as dominant drivers of thyroid carcinoma, jointly affecting approximately 70% of the cases^36,42–46^. RAS and BRAF are generally recognized as the main driver mutations in FTC and PTC, respectively, which predominantly change protein activity and do not necessarily associate with altered expression levels, whereas RET translocations can lead to changes in both protein activity and expression levels, depending on the type of alteration. In this dataset, however, we observed that NRAS (isoform of RAS) was slightly upregulated (FDR < 0.05, Log_2_(FC) ≥ 1) in FTC (FDR = 4.66×10^−2^, Log_2_(FC) = 1.30) as well as PTC (FDR = 1.35×10^−2^, Log_2_(FC) = 1.08) samples when compared to the healthy thyroid tissues, while the RAS Q61R mutation was detected in a single FA and PDTC (12.5%) sample by IHC staining, each (**Table S1**). Upon RAS pathway activation, KSR1, a scaffold protein within the MAPK signaling pathway, can be dephosphorylated at the residue S406, leading to the dissociation of 14-3-3 signaling adaptor proteins^47,48^. This leads to the translocation to the plasma membrane, where MAP2K1 and MAP2K2 get phosphorylated and activated. Interestingly, most of the investigated thyroid tumors showed reduced KSR1 S406 phosphorylation levels (FA: FDR = 4.60×10^−2^, Log_2_(FC) = –1.39, FTC: FDR = 3.40×10^−2^, Log_2_(FC) = –1.43, PTC: FDR = 2.96×10^−3^, Log_2_(FC) = –1.55, PDTC: FDR = 4.86×10^−3^, Log_2_(FC) = –1.75), and in FA (FDR = 2.64×10^−2^, Log_2_(FC) = 1.55) and in FTC additionally the MAP2K2 (FDR = 2.74×10^−2^, Log_2_(FC) = 1.24) kinase was overexpressed. We further noticed a downregulated (FDR < 0.05, Log_2_(FC) ≤ –1) phosphorylation of the BRAF S729 residue (FDR = 1.59×10^−2^, Log_2_(FC) = –1.09) in PTC that, when phosphorylated, mediates an inhibitory effect on its signaling activity due to 14-3-3 protein binding^49^. Apart from the phosphorylation status of BRAF, we further detected positive BRAF V600E IHC staining exclusively in PTC with a high prevalence of 69% (11/16), which activates BRAF and downstream MAPK signaling^50^. With regards to RET mutated medullary thyroid carcinoma (MTC), it has previously been shown that the tyrosine-protein kinase CSK mediates cellular proliferation via ERK and AKT signaling in human cancer cells^51^. Here, we observed enhanced CSK protein expression (FDR = 3.45×10^−3^, Log_2_(FC) = 1.41) as well as upregulation of the AKT1 target phosphosite S143 in the DNMT1 protein in PTC (FDR = 1.22×10^−2^, Log_2_(FC) = 1.07). Independently, AKT1 phosphorylation at the S126 residue, which was suggested to induce the AKT1 enzymatic activity^52^, was upregulated in PDTC (FDR = 2.34×10^−2^, Log_2_(FC) = 1.31). Interestingly and although no direct link to its activity is known, the S122 residue was among the most upregulated phosphorylation sites in FA samples (FDR = 2.78×10^−4^, Log_2_(FC) = 3.34). The STAT1 transcriptional activity has previously been described to be enhanced in PTC cells harboring RET/PTC gene rearrangement^53^. In this study, we also observed a high overexpression of the STAT1 protein in PTC^26^ samples (PTC: FDR = 1.11×10^−2^, Log_2_(FC) = 2.37). The role of STAT1 in cancer, e.g. transcriptional regulation of genes involved in apoptosis and cell cycle, is diverse and appears to be context-specific^54^. Finally, we observed upregulation of the MAPK14 T180 phosphoresidue, a site important for its kinetic activity, in both benign adenoma and well-differentiated carcinoma samples (FA: FDR = 4.16×10^−2^, Log_2_(FC) = 1.15, OA: FDR = 3.84×10^−2^, Log_2_(FC) = 1.55, FTC: FDR = 1.05×10^−2^, Log_2_(FC) = 2.33, PTC: FDR = 4.45×10^−3^, Log_2_(FC) = 1.66) as well as the additional upregulation of its Y182 phosphoresidue (FA: FDR = 3.18×10^−2^, Log_2_(FC) = 1.69, OA: FDR = 3.57×10^−2^, Log_2_(FC) = 1.52, FTC: FDR = 1.53×10^−2^, Log_2_(FC) = 2.31, PTC: FDR = 4.25×10^−3^, Log_2_(FC) = 1.72).

Furthermore, we found upregulated phosphosites in the Bcl2-associated agonist of cell death (BAD) protein, which has a role in inhibiting the apoptotic function of BAX and BAK proteins^55^, in FA (S118 FDR = 4.26×10^−2^, Log_2_(FC) = 1.78), OA (S99 FDR = 4.52×10^−2^, Log_2_(FC) = 2.00), and PDTC (S99 FDR = 1.95×10^−2^, Log_2_(FC) = 1.84) samples. The S99 anti-apoptotic BAD phosphorylation site can be phosphorylated by AKT1^56^ and PAK1^57^ kinases. BAX itself has in previous studies been found to be overexpressed in more aggressive forms of thyroid cancer^58^. Here, we observed upregulated expression levels of BAX in PTC (FDR = 2.67×10^−2^, Log_2_(FC) = 1.28) and benign FA (3.69×10^−2^, Log_2_(FC) = 1.74), but not in other tumor subtypes.

In line with previous observations that linked mutations of the AKT/MTOR pathway, including MTOR itself, more strongly to the PDTC than PTC^16^ subtype, we found differential expression of MTOR exclusively in PDTC samples (FDR = 9.57×10^−3^, Log_2_(FC) = 1.50). We additionally detected upregulation of several of its proline-rich AKT1 substrate 1 (AKT1S1) phosphorylation sites (**Table S3**). Among them, the mTORC1 signaling activating residue S203^59^, was found phosphorylated more strongly in all thyroid tumors investigated (FA: FDR = 3.92×10^−3^, Log_2_(FC) = 1.79, OA: FDR = 1.66×10^−2^, Log_2_(FC) = 1.50, FTC: FDR = 3.04×10^−2^, Log_2_(FC) = 1.28, PTC: FDR = 4.68×10^−4^, Log_2_(FC) = 1.44, PDTC: FDR = 4.94×10^−3^, Log_2_(FC) = 1.24). In addition, we also observed overexpression of the ANP32E epigenetic regulator in PTC (FDR = 6.25×10^−3^, Log_2_(FC) = 1.27), a protein known to promote PTC cell proliferation, growth, and migration through AKT/MTOR signaling^60^.

Overall, the complex molecular adaptations show tumor subtype-specific alterations in several cancer drivers, protein kinases, TFs, and EFs. Particularly prominent changes were observed along the well-established oncogenic RAS/BRAF/MAPK and AKT/MTOR signaling axes in malignant subtypes.

### Functional analysis of proteomic profiles reveals distinct and overlapping molecular pathways in different thyroid tumors

To assess cellular functional roles affected by the observed proteomic alterations across the studied subtypes of thyroid tumors, we performed gene set enrichment analysis (GSEA) using the MSigDB hallmark gene set annotations. For this, all measured proteins in the individual subtypes were ranked in accordance to their expression and significance values in comparison to healthy thyroid tissues (**Table S4**). With GSEA the resulting enrichment score (ES) indicates the extent to which a particular hallmark gene set is enriched among the top or bottom ranked proteins in the analyzed set. The normalized enrichment score (NES) further accounts for variations in the gene set size of annotated pathways^61^. We found that 16 hallmark gene sets had statistically significant NES (FDR < 0.05) in at least one studied tumor subtype (**Fig. 3A-B**). The majority, i.e. 11 of the gene sets were significantly enriched only in a single thyroid tumor subtype thus indicating possible functional differences among all subtypes. For instance, PTC samples were associated with a strong inflammatory response (*Interferon Alpha/Gamma Response*) that was previously suggested to be linked with an unfavorable prognosis^62^, and only benign adenomas (both FA and OA) had higher expression levels of proteins with a role in *Oxidative Phosphorylation*, i.e. metabolic route that is usually downregulated in more aggressive tumors^63^. Additionally, we found a negative enrichment of the *Epithelial Mesenchymal Transition* gene set only in OA. On the other hand, clear oncogenic signatures were evident within the PDTC tumor subtype, i.e. the most aggressive tumor form studied here. This included the enrichment of two gene sets associated with cell cycle progression and proliferation (*E2F Targets*, *G2-M Checkpoint*) as well as an enrichment of MYC targets (*Myc Targets V1*), whose upregulation was previously shown to contribute to the dedifferentiation of follicular cell derived thyroid cancer^22,64^. The five top ranked proteins among the MYC target pathway included PPM1G (FDR = 2.29×10^−5^, Log_2_(FC) = 3.79), PSMD8 (FDR = 5.04×10^−3^, Log_2_(FC) = 1.57), ERH (FDR = 1.15×10^−2^, Log_2_(FC) = 1.85), EIF3D (FDR = 1.90×10^−2^, Log_2_(FC) = 1.94), and SRM (FDR = 8.40×10^−3^, Log_2_(FC) = 1.50), all of which have been previously associated with progression and poor prognosis in different cancer types^65–69^.

**Figure 3:**
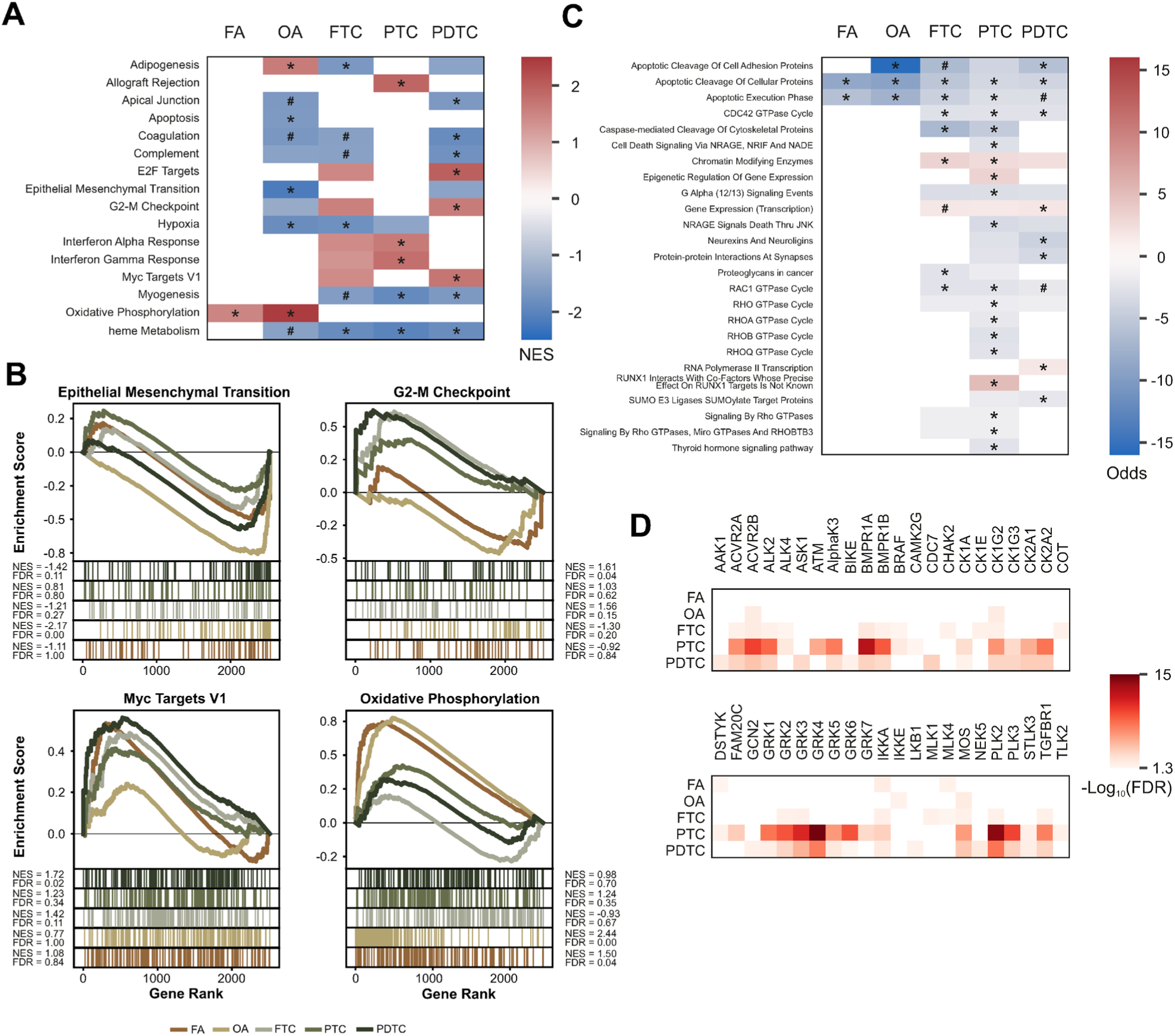
Pathway and kinase analysis of thyroid neoplasms of follicular origin. (**A**) Heatmap of enriched pathways in thyroid neoplasms compared to healthy controls using GSEA (hallmark gene sets) based on proteomic measurements. Entries with at least one FDR < 0.05 are shown with * indicating FDR < 0.05, # FDR < 0.1, and colored cells FDR < 0.25. The color scheme illustrates the associated NES values. (**B**) GSEA enrichment plots of representative pathways. (**C**) Heatmap of over-represented pathways in thyroid neoplasms compared to healthy controls using ORA (KEGG and Reactome gene sets) based on phosphoproteomic measurements. Entries with at least one FDR < 0.05 are shown with * indicating FDR < 0.05, # FDR < 0.1, and colored cells FDR < 0.25. The color scheme illustrates the associated Odds ratios. (**D**) Significant (FDR < 0.05) predictions of enhanced kinase activities in thyroid neoplasms compared to healthy controls using The Kinase Library tool^71^ from PhosphoSitePlus^38^. The color scheme illustrates the associated FDR values.

Further, we assessed if proteins whose phosphosites were significantly up-or downregulated in tumor samples (FDR < 0.05, |Log_2_(FC) ≥ 1|), and for which deregulation could not simply be explained by changes in protein expression levels (see Methods), were over-represented in specific biological pathways. For this, we applied over-representation analyses (ORA) by using KEGG^37^ and Reactome^70^ signaling pathway annotations and compared significantly deregulated phosphoproteins to all measured (phospho-)proteins (**Fig. 3C**, **Table S5**). The ORA resulted in the identification of 25 significantly (FDR < 0.05) over-represented pathways, and included both shared and subtype-specific functions. For instance, phosphorylation levels of proteins involved in the apoptotic pathways (*Apoptotic Cleavage Of Cellular Proteins*, *Apoptotic Execution Phase*) were downregulated in all thyroid tumors, indicating evasion from programmed cell death via apoptosis. Exclusively in PTC samples, the downregulated death signal pathways were further found to involve the protein NRAGE (*Cell Death Signaling Via NRAGE, NRIF And NADE*, *NRAGE Signals Death Thru JNK*). The phosphorylation levels of two epigenetic pathways (*Chromatin Modifying Enzymes*, *Epigenetic Regulation Of Gene Expression*) were found upregulated in both studied well-differentiated carcinoma samples (FTC, PTC). Those of the gene expression machinery (*Gene Expression (Transcription)*, *RNA Polymerase II Transcription*), however, were solely over-represented in PDTC thyroid samples. Among others, epigenetic pathways upregulated in both FTC and PTC shared enhanced phosphorylation levels of histone deacetylase complex subunits (SAP30, SAP30L) and transcriptional repressors (GATAD2A, GATAD2B). Further instances of proteins that had upregulated phosphorylation levels of chromatin modifying enzymes in both FTC and PTC samples included histone-lysine N-methyltransferases (SETD1B, SETD2) and histone acetyltransferases (KAT5, KAT7). The epigenetic regulation pathway in PTC additionally included upregulation of the DNMT1 S143 phosphoresidue, which is a direct target of the AKT1 kinase that itself is known to inhibit anti-tumorigenic mechanisms in thyroid tumors^17^. Collectively, the over-represented pathways reflect profound changes in the molecular profiles of thyroid tumor.

In order to specifically predict enhanced kinase activities based on the known activity footprints of individual kinases, we made use of The Kinase Library tool^71^ (see Methods), which is built based on kinase preferences observed from a systematic screen of synthetic peptides. For this, we assessed quantitative levels of the phosphopeptides with the measured phosphosites in the center (up to 15 amino acids in the direction of both N and C terminus, when possible) in the individual thyroid tumors when compared to healthy tissue. As expected from the overall lower fraction of differentially regulated phosphosites in both adenoma samples (FA, OA, **Fig. 2B**), these subtypes had the lowest number of predicted upregulated protein kinases (**Fig. 3D**, **Table S6**). Furthermore, this showed that the BRAF signaling was not only prominently upregulated in the well-differentiated PTC (FDR = 6.19×10^−3^) samples, which are associated with a high prevalence of activating BRAF V600E mutations (69%), but also in FTC (FDR = 3.86×10^−2^). In addition, the serine-protein kinase ATM was predicted to be highly active in PTC (FDR = 4.15×10^−6^). The standard mechanism of ATM activation is initiated by DNA double strand breaks^72^ and its inferred overactivation is in line with a previous study, which found prominent upregulation of ATM signaling in PTC based on transcriptomic data^73^. In addition, we observed enhanced activities of all seven G protein-coupled receptor (GPCR) kinase family members (GRK1-7) in PTC samples (FDR < 0.05). The GRK2-3 kinases were also significant in the FTC subtype and GRK1-6 in PDTC (**Fig. 3D**). It is worth noting that the prediction tool might struggle with distinguishing activities of the kinase members of the same family as phosphomotifs they recognize share considerable similarities. GRKs exhibit diverse functional roles in a variety of cancer types by modulating GPCR and growth factor receptor signaling^74^. GRK2 is able to inhibit the activity of the TSH receptor (TSHR), whereas GRK5 can inhibit its desensitization^75^. Activation of TSHR is known to increase cancer cell proliferation in differentiated thyroid carcinoma (FTC, PTC)^75^. Furthermore, GRK6 has been shown to exhibit oncogenic activity and to correlate with poor prognosis in PTC^76^. Additionally, Polo-like kinases (PLKs) were predicted to have upregulated activities in more aggressive subtypes: FTC, PTC, and PDTC all showed footprints of increased activity of PLK2 (FDR < 0.03), and the latter two tumor subtypes were additionally connected with enhanced PLK3 activity (FDR < 0.01). PLKs have a wide range of cellular functions that include cell cycle progression and survival^77^. Volasertib is a potent inhibitor of PLK1-3^78^ and it has previously been shown that treatment with Volasertib inhibits cell proliferation in FTC, PTC, and ATC cell lines^79,80^.

Taken together, the molecular changes in the various thyroid tumors revealed both subtype-specific and shared biological processes. Enhanced BRAF (in PTC and, contrary to expectations, also in FTC), GRK5 and GRK6 (both in PTC and PDTC), as well as PLK2 (in FTC, PTC, and PDTC) and PLK3 (PTC, PDTC) kinase activities are observed in different malignant thyroid tumors indicating possible shared molecular mechanisms in the development of more aggressive thyroid tumors. Of interest is also the inferred increase in ATM activity in PTC, which might reflect a different aberrant process specifically present in this tumor type.

### FA and FTC proteomes display detectable molecular adaptations during disease progression despite their relative proximity

FTC can arise from FA and it has been hypothesized that the potential progression could be explained through the accumulation of multiple successive minor alterations rather than through a few major driver mutations. Moreover, the current recommendation is that the distinction between the two subtypes should be based on morphological differences in invasiveness, i.e. capsular and most importantly vascular invasion, rather than on molecular signatures^3,81^. In support of this, recent studies have shown that the global proteome analysis of dozens of FA and FTC samples was not able to clearly distinguish the two subtypes^26,31^. Given that integrated phosphoproteomics analysis has previously contributed to the characterization of stratified molecular subtypes within PTC^27^ and recurrence risk groups within MTC^82^ samples, and the fact that FA and FTC samples were clearly separated in our cluster analyses (**Fig. 1B-E**), we have extended the comparative proteomic and phosphoproteomic analyses to FA and FTC to more closely assess the extent of the molecular differences.

The PCA analysis (**Fig. 4A-B**) restricted to FTC and FA subtypes successfully clustered the samples according to their group identity, which is not possible based on transcriptomics or genomics ^83–85^. Therefore, we performed a DEA comparing the protein and phosphopeptide expression levels between the FA and FTC samples. We found that in total 81 proteins and 229 phosphopeptides had significantly higher expression levels in FTC, whereas 22 proteins and 225 phosphopeptides were measured with higher expression levels in FA (moderated *t*-test, FDR < 0.05 and absolute Log_2_(FC) ≥ 1, **Fig. 4C-D**, **Table S2**-**S3**). These numbers are markedly lower than when comparing expression levels of FA and FTC with healthy tissue controls, thus demonstrating the close similarity of the two subtypes at the molecular level. Among others, this highlighted increased expression of TSHR (FDR = 3.98×10^−2^, Log_2_(FC) = 1.18) in FTC compared to FA. TSHR is important for cell growth of differentiated thyroid carcinoma like FTC and has been demonstrated as useful target of chimeric antigen receptor (CAR) cell therapies in the past^75,86^. Furthermore, FTC samples had an enhanced expression of the ubiquitin hydrolase USP7 (FDR = 3.62×10^−2^, Log_2_(FC) = 1.03), which promotes PTC proliferation when overexpressed^87^.

**Figure 4:**
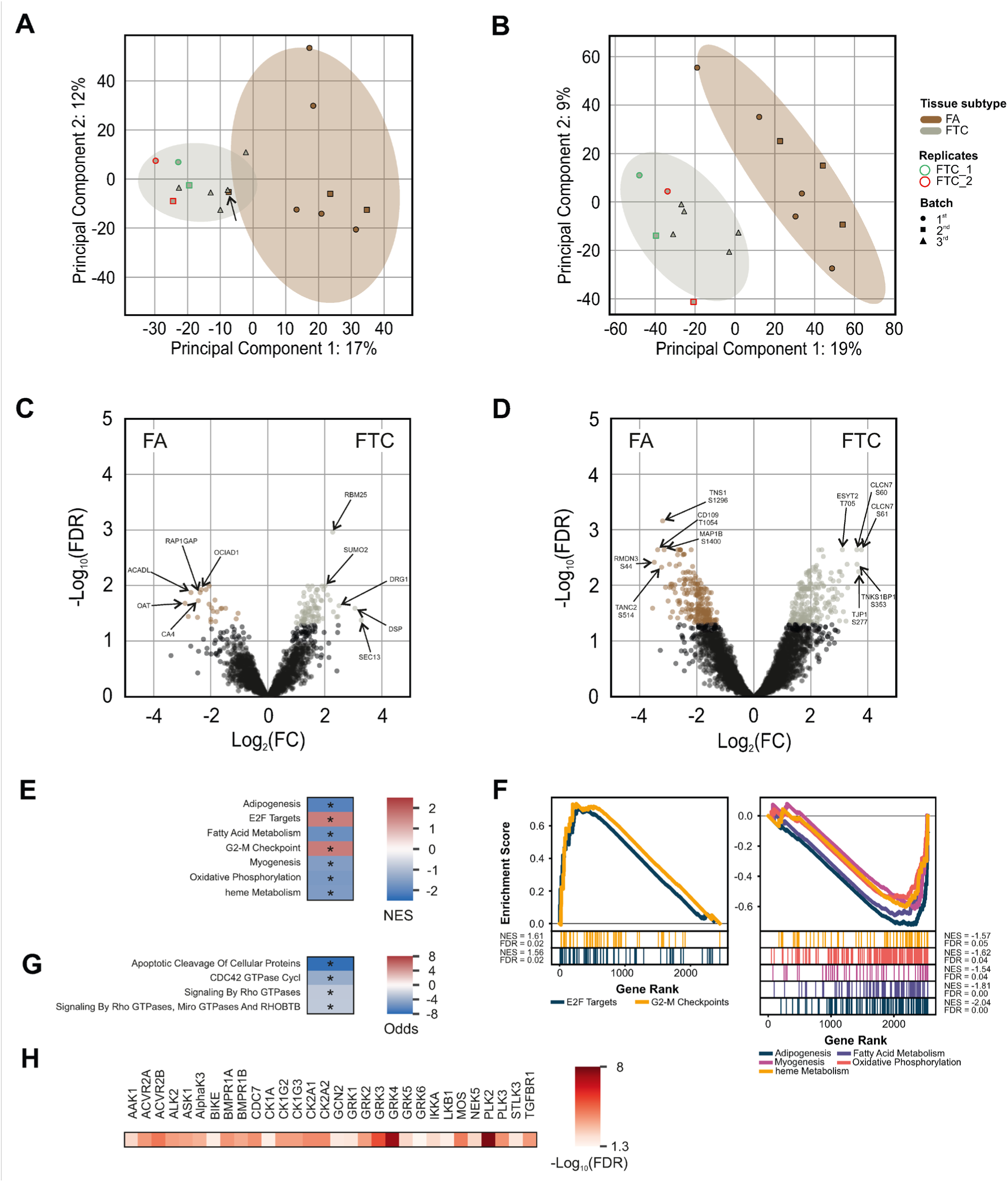
Molecular differences between follicular adenoma and carcinoma. **A, B** Principal component analysis (PCA) clustering including 95% confidence ellipses based on the global (**A**) proteome and (**B**) phosphoproteome of FA and FTC samples. **C, D** Volcano plots of quantified (**C**) proteins and (**D**) phosphopeptides comparing FTC with FA. Colored entries represent significant hits with FDR < 0.05 and absolute Log_2_(FC) ≥ 1. Top five up– and downregulated hits were annotated based on ranking according to –Log_10_FDR × Log_2_(FC). (**E**) Heatmap of enriched pathways in thyroid neoplasms compared to healthy controls using GSEA (hallmark gene sets) based on proteomic measurements. Entries with FDR < 0.05 values (highlighted with *) are shown. The color scheme illustrates the associated NES values. (**F**) GSEA enrichment plots of all pathways. (**G**) Heatmap of over-represented pathways in FTC compared to FA using ORA (KEGG and Reactome gene sets) based on phosphoproteomic measurements. Entries with FDR < 0.05 values (highlighted with *) are shown. The color scheme illustrates the associated Odds ratios. (**H**) Significant (FDR < 0.05) predictions of enhanced kinase activities in FTC compared to FA, using The Kinase Library tool^71^ from PhosphoSitePlus^38^. The color scheme illustrates the associated FDR values.

We used the list of all measured proteins ranked according to the significance in the difference of their expression levels between the FA and FTC samples for GSEA based on the MSigDB hallmark gene sets (**Fig. 4E-F**, **Table S4**). This revealed an enrichment in the cell cycle related targets of the E2F transcription factors (*E2F Targets*) in FTC. E2F1 has previously been suggested as a transcription factor that potentially underlies gene expression differences between FTC and FA^88^. Moreover, the FTC proteome was enriched in the *G2-M Checkpoint* pathway, which is also related to cell cycle progression. The upregulated cell cycle in FTC might reflect rapid and uncontrolled cell proliferation, which leads to tumor capsular penetration and invasion by the cells, a fundamental morphological feature that differentiates FTC from FA^3^. In contrast, FA was mainly linked with higher levels of several metabolic (*Fatty Acid Metabolism*, *heme Metabolism*) and tissue developmental (*Adipogenesis*, *Myogenesis*) pathways. ORA of the differentially regulated phosphopeptides in KEGG^37^ and Reactome^70^ pathways highlighted the downregulation of *Apoptotic Cleavage Of Cellular Proteins* and GTPase related pathways (*CDC42 GTPase Cycl*, *Signaling By Rho GTPases*, *Signaling By Rho GTPases, Miro GTPases AND RHOBTB*) in FTC (**Fig. 4G**, **Table S5**). As the same pathways were previously identified as hallmarks of FTC compared to healthy tissue, this observation is consistent with the expected similarity of FA with healthy tissue.

By applying The Kinase Library tool^71^ (see Methods), we inferred a total of 30 kinases with enhanced activities in FTC compared to FA (**Fig. 4H**, **Table S6**). Of these, 12 kinases have already been identified in comparison to healthy tissue, indicating tumorigenic functions not only in the development of FTC but also progression from FA to FTC. Among the newly identified 18 kinases, 11 were identified in this study to be associated with PTC and PDTC, and the remaining 7 kinases (AAK1, ASK1, BIKE, CDC7, GCN2, LKB1, and NEK5) were inferred as upregulated in PDTC, the most aggressive thyroid malignancy studied here. Already identified entries include GRK2 (FDR = 3.06×10^−4^) and PLK2 (FDR = 1.95×10^−8^) with effects on TSHR activity^75^ and cell cycle progression^77^, respectively. GRK5 (FDR = 4.94 ×10^−3^), GRK6 (FDR = 4.96×10^−2^), and PLK3 (FDR = 1.73×10^−4^) were also found to be associated with increased activity in FTC compared to FA, suggesting a role in the malignancy of FTC.

Collectively, the low number of significantly differentially regulated entries shows relatively close proteomic and phosphoproteomic profiles of FA and FTC. The data analysis, however, revealed that the malignancy of FTC in comparison to benign FA is associated with an enhanced TSHR expression, aberrant kinase activities, as well as changes in biological pathways related to cell cycle, metabolism, apoptosis, and tissue development.

In **Fig. 4A**, the black arrow points out a sample (FA_8) diagnosed as FA while it is clustered within the FTC ellipse and almost directly overlaps with an FTC case (FTC_3). Despite the high variability in the ellipses, this overlap raised suspicion. To further investigate, we subjected the FFPE (Formalin-Fixed, Paraffin-Embedded) blocks of the corresponding sample to X-ray virtual histology, as described previously^3,89^. The virtual histology revealed two instances of vascular invasion (VI) that were missed by conventional histological analysis. Following this, the relevant FFPE blocks were serially sectioned, and whole slide images were examined to correlate the virtual histology findings with traditional histology, as shown in **Fig. 5**. The serial sectioning confirmed the VIs identified in the virtual histology. Consequently, the FA diagnosis was revised and thus referred to as thyroid with uncertain malignant potential. The proteomic clustering previously thought to be an outlier within the FA group was, in fact, correctly reflected by its uncertain malignant potential.

**Figure 5:**
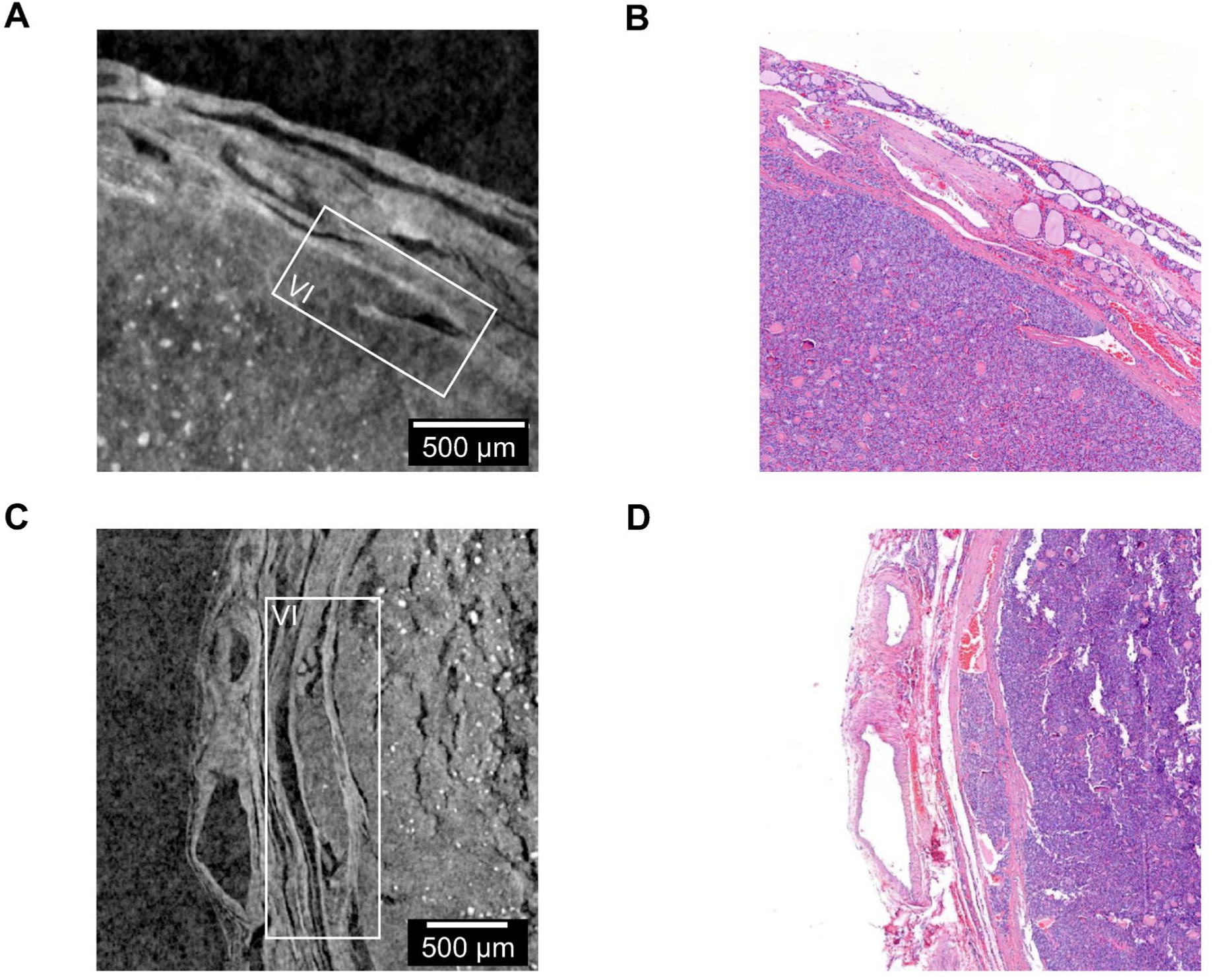
Histology of the thyroid with uncertain malignant potential. **A** and **C** show the VIs identified by X-ray virtual histology inside the white rectangles, while **B** and **D** present the corresponding conventional histology images, which confirm the virtual histology observations.

## Discussion

In this study, we conducted an integrative proteomics and phosphoproteomics analysis across different thyroid tumors of follicular origin, whose precise clinical classification remains a challenge to this date^3^. Phosphoproteomics has previously only been applied to the PTC^27^ follicular subtype. Therefore, the new level of tumor characterization offered us the opportunity to not only compare molecular adaptations at the level of protein expression but also to infer more directly upregulated signaling pathways and kinase activities among various thyroid tumor subtypes of the same cellular origin (FA, OA, FTC, PTC, PDTC). Besides the unbiased and global expansion of our molecular understanding, we particularly focused on reconciling the adaptations in the well-established oncogenic RAS/BRAF/MAPK and AKT/MTOR signaling axes with existing knowledge. The applied analysis protocol, including imputation and batch correction following concepts of PhosR^30^ and MSstatsPTM^90^, was able to preserve biologically meaningful differences in expression data despite complex experimental design. As a corollary, clustering approaches successfully stratified malignant (FTC, PTC, PDTC) and healthy thyroid tissues. Also, the proteomic and phosphoproteomic quantitative measurements successfully stratified FA and FTC subtypes, which was not possible in previous studies^26,31^. Interestingly, an identified outlier in the PCA of the proteomics data, i.e. a specific FA sample co-clustering with the FTC group, could be verified as a misdiagnosed case with uncertain malignant potential by X-ray virtual histology. Along the axis of malignancy we observed the importance of key biological and tumor-associated cellular programs including regulation of cell cycle control, metabolic reprogramming, apoptosis, tissue development, and specific signaling activities.

By comparing the molecular profiles of the thyroid tumors of follicular origin individually with healthy control tissues, we identified multiple shared and subtype-specific changes in protein expression and phosphorylation levels. As expected, benign thyroid tumors (FA, OA) showed less profound changes in dysregulation of downstream biological pathways and inferred changes in kinase activities. However, OA deviated from the expected pattern with a surprisingly high number of differentially expressed proteins, which is likely underlined by compensatory mitochondrial biogenesis. Many of the molecular changes identified in this study were found within the RAS/BRAF/MAPK and AKT/MTOR signaling axes, especially in malignant tumors. This is illustrated with the upregulated expression of NRAS protein and predicted enhanced BRAF kinase activity in well-differentiated FTC and PTC as well as the overexpression of MTOR in PDTC. In general, changes in these signaling pathways were highly expected. However, the increased NRAS expression in the PTC group, in which most samples showed positive BRAF V600E IHC staining (69%), was unexpected and could indicate an increased RAS signal independent of a RAS Q61R mutation. Furthermore, and also contrary to the prevailing paradigm, the predicted increased signaling activity of BRAF occurred not only in PTC but also in FTC.

Additionally, analyses of phosphoproteomic profiles highlighted increased activity of the ATM kinase in PTC. The fact that several ATM inhibitors, as monotherapies or in combination with DNA-damaging agents, have entered clinical trials for the treatment of solid tumors^91^ suggest that ATM may also be of interest as a therapeutic target for the treatment of severe PTC cases. Besides, this study pointed to the high activity of the PLK2 and PLK3 as well as GRK5 and GRK6 kinases in malignant tumors, in agreement with the previous notions of their roles in the development and progression of thyroid tumors.

In summary, our integrative approach provides a broad mapping of the molecular landscapes across various thyroid tumors of follicular origin. As a result, we found alterations in protein expression and phosphorylations profiles pointing towards changed molecular regulations within several important biological pathways, including cell cycle, apoptosis, and metabolic reprogramming. This study showed that activation of RAS/BRAF/MAPK and AKT/MTOR signaling axes plays a role in the development of various and especially malignant thyroid cancer subtypes, and that upregulation of several kinases (e.g. GRK5, GRK6, PLK2, and PLK3) contributes to more aggressive phenotypes. However, the observation that both RAS and BRAF kinases play a role in FTC as well as PTC at the level of the proteome and phosphoproteome, respectively, contradicts the prevailing paradigm and therefore warrants further functional investigation. The generated data provides a comprehensive set for integration with complementary omics datasets with the potential to improve the classification scheme of thyroid tumors of follicular origin and identify new therapeutically targetable disease drivers.

## Materials and methods

### Thyroid samples

The Tissue Bank Bern (TBB) of the University of Bern and the tissue biobank of the University Hospital of Zurich manage tissue samples and patient information according to respective regulations under fulfillment of the Swiss Biobanking Platform requirements. The Institute of Tissue Medicine and Pathology of the University of Bern and the Institute of Pathology and Molecular Pathology of the University Hospital of Zurich provided the samples according to the protocols approved by the cantonal ethics commissions (KEK BE 2018-01657 and KEK ZH 2016-01575). Biobanked resected treatment naïve thyroid tissues were reviewed, classified according to WHO guidelines, and selected by O. S., A. P., and A. W., all experienced pathologists. Fresh frozen and OCT embedded tissues were shipped on dry ice to the Functional Genomics Center Zurich (FGCZ) core facility for measurements. A formalin fixed paraffin embedded representative block from every sample was biopsied with 0.6 mm core to obtain both tumor and normal tissue to produce a Tissue MicroArray Block of all samples, which was subsequently stained with an anti-BRAF V600E monoclonal antibody (clone: VE1, dilution: ready to use; Roche Diagnostics) using Benchmark Ultra System (Ventana Roche Switzerland), and an anti-NRAS Q61R monoclonal antibody (clone: SP174, dilution: 1:50; Abcam) using the Leica Bond III system (Biosystems Switzerland), according to the manufacturer’s recommendations. Strong cytoplasmic staining for BRAF V600E with on-slide positive control as well as diffuse moderate to strong membranous and cytoplasmic staining for RAS Q61R were considered positive (**Table S1**).

### Protein extraction and digestion

The quantitative proteomics and phosphoproteomics measurements were performed by the FGCZ. Samples embedded in OCT were washed with 100 µL of 100% ethanol followed by a washing step with 200 µL ddH_2_O. To each sample 100-200 µL of lysis buffer (4% SDS, 100 mM Tris / HCL pH 8.2) were added. Protein extraction was carried out using a tissue homogenizer (TissueLyser II, QIAGEN) by applying 2× 2 min cycles at 30 Hz followed by boiling at 95 °C for 10 min and treating with High Intensity Focused Ultrasound (HIFU) for 1 min setting the ultrasonic amplitude to 100%. Insoluble debris were collected by centrifugation at 20’000× g for 10 min. Protein concentration was determined using the Lunatic UV/Vis polychromatic spectrophotometer (Unchained Labs). Equal total protein amounts were taken from each sample, reduced and alkylated with 5 mM TCEP (tris(2-carboxyethyl)phosphine) and 15 mM chloroacetamide at 30 °C for 30 min in the dark.

SP3-based protein purification, digestion and peptide clean-up were performed using a KingFisher Flex System (ThermoFisher Scientific) and Carboxylate-Modified Magnetic Particles (GE Life Sciences; GE65152105050250, GE45152105050250)^92,93^. Samples were diluted with 100% ethanol to a final concentration of 60% ethanol. After loading beads, wash solutions and samples in 96 deep-well or micro-plates were transferred to the KingFisher system. The following steps were carried out on the robot: collection of beads from the last wash, protein binding to beads, washing of beads in wash solutions (80% ethanol) and addition of digestion solution for overnight offline digestion on a Thermomixer at 37 °C with a trypsin:protein ratio of 1:50 in 50 mM Triethylammoniumbicarbonat (TEAB). After the digest, beads were again transferred to the KingFisher to collect the digest solution and subsequent water elution, which were subsequently combined and dried to completeness until further processing.

### Phosphopeptide enrichment

Fe-NTA beads (Cube Biotech) and a KingFisher Flex System (ThermoFisher Scientific) were used for the enrichment of phosphopeptides according to a by Leutert *et al*. published protocol^93^. The samples were dissolved in 200 µL loading buffer (80% ACN, 5% TFA) before a volume corresponding to 2 µg total protein was removed for global proteome analysis. Enrichment of phosphopeptides was carried out in the following steps: washing of the magnetic beads in loading buffer (5 min), binding of the phosphopeptides to beads (20 min), 3× washing in loading buffer and eluting the phosphopeptides from the magnetic beads (1 M NH4OH, 10 min). The phosphopeptides were dried to completeness and re-solubilized in 10 µL of 3% acetonitrile, 0.1% formic acid for subsequent MS analysis.

### HPLC-MS/MS analysis

Phosphoproteomics sample were analyzed on an Orbitrap Fusion Lumos (Thermo Scientific) equipped with a Digital PicoView source (New Objective) and coupled to an M-Class UPLC (Waters). Solvent composition of the two channels was 0.1% formic acid for channel A and 99.9% acetonitrile in 0.1% formic acid for channel B. For each sample 4 μL of peptides were loaded on a commercial ACQUITY UPLC M-Class Symmetry C18 Trap Column (100 Å, 5 µm, 180 µm × 20 mm, Waters) connected in-line to an ACQUITY UPLC M-Class HSS T3 Column (100 Å, 1.8 µm, 75 µm × 250 mm, Waters). Peptide separation was carried out at a constant flow rate of 300 nL×min^−1^. After a 3 min initial hold at 5% B, a gradient from 5 to 22% B in 83 min and from 22 to 32% B in an additional 10 min was applied. Samples were acquired in randomized order with the following DDA acquisition parameters: full-scan MS spectra (375−1500 m/z) were acquired at a resolution of 120’000 at 200 m/z after accumulation to an automated gain control (AGC) target value of 400’000 or 40 ms inject time. Precursors with an intensity above 5’000 were selected for MS/MS, where ions were isolated using a quadrupole mass filter with 1 m/z isolation window and fragmented by higher-energy collisional dissociation (HCD) using a normalized collision energy of 35. Fragments were detected in the linear ion trap using adapted “Universal Method” settings: the scan rate was set to rapid, the automatic gain control was set to 10’000 ions, and the maximum injection time was 120 ms. Charge state screening was enabled, and singly, unassigned charge states and charge states higher than seven were excluded. Precursor masses previously selected for MS/MS measurement were also excluded from further selection for 20 s.

MS analysis of the proteome samples was performed on the same system. For each sample, 10% the of peptides were loaded and analyzed with the same acquisition parameters as described above except for ion selection in the quadrupole mass filter, which was done with a 0.8 m/z isolation window and a maximum MS2 ion injection time of 50 ms. Both the proteome and phosphoproteomics samples were acquired using internal lock mass calibration on m/z 371.1010 Th and 445.1200 Th. The MS data were handled using the local laboratory information management system (LIMS)^94^.

### Protein and phosphopeptide identification and LFQ

After acquisition, MS data were processed by MaxQuant^28,29^ version 2.0.1.0, whereas the integrated Andromeda search engine was used for protein identification. The spectra were searched against a canonical proteome reference of Homo sapiens (UniProt^41^: UP000005640, downloaded 2023-03-30), concatenated to common protein contaminants. The carbamidomethylation of cysteine was set as fixed modification and methionine oxidation as well as N-terminal protein acetylation were set as variable. For the phosphoproteome search, serine, threonine and tyrosine phosphorylation were additionally set as variable modifications. Enzyme specificity was set to trypsin/P, the minimal peptide length to 7 amino acids and the maximum of missed-cleavages to two. MaxQuant default search settings were used. The maximum FDR was set to 0.01 for peptides and 0.05 for proteins. LFQ was enabled applying a 2 min window for match between runs. Each file is kept separate in the experimental design of MaxQuant to obtain individual quantitative values.

### Data handling

The data analysis was based on quantitative matrices of protein intensities (proteinGroups.txt) and individual peptide phosphorylation intensities (Phospho_STY_Sites.txt). Commonly occurring contaminants (fasta file provided by MaxQuant, e.g. keratins and trypsin), entries matching the reversed part of the decoy database, and proteins only identified by peptides carrying modified amino acids were excluded from the subsequent analysis. Proteomic entries that were only identified by a single peptide were not considered. Furthermore, phosphopeptides were required to have a localization confidence above 75% and a posterior error probability below 1%. All entries were filtered for those with measured values in at least 50% of the replicates in at least one of the conditions investigated.

The LFQ intensities were Log_2_ transformed. Missing values were imputed separately for each replicate within a sample group following concepts of the PhosR method^30^. Here, missing values that are consistently absent in a certain group and values missing only in a small fraction of samples are distinguished. For this, two different normal distributions shifted left from the mean of the measured values were constructed for each replicate by using the PaDuA library^95^, of which missing data entries were randomly sampled from either of the distributions depending on the fraction of measured values in the corresponding condition. When more than 50% of the replicates in a sample group were measured, the values were sampled from a distribution shifted left 0.5 of the standard deviation (SD) of the original mean. Otherwise, the negative shift was 1.8 SD. In both cases, a width of 0.3 SD was applied. LFQ intensities were subsequently median-centered across each replicate. Batch correction was applied using the pyComBat^96^ tool.

### Data analysis

For HC the seaborn^97^ library with z-scored row entries was used. For PCA the scikit-learn^98^ was implemented. DEA on the level of proteins and phosphopeptides was conducted based on moderated *t*-tests in limma^99^. Biological replicates were not considered in the analyses. Obtained *p*-values were corrected for multiple testing using the Benjamini-Hochberg (BH) method^100^. Implemented thresholds for significance included FDR < 0.05 and an absolute Log2(FC) ≥ 1. GSEA of proteomics data within MSigDB hallmark gene sets was conducted with GSEApy^101^, applying a gene ranking metrics according to Log_2_(FC) × –Log_10_(FDR). Resulting *p*-values were again corrected using the BH^100^ method implementing a threshold of FDR < 0.05. ORA and prediction of enhanced kinase activities were based on phosphoproteomics data. Therefore, an additional threshold was applied to ensure that changes of phosphorylation levels were not observed simply due to altered protein expression. Following concepts of the MSstatsPTM^90^ approach, we implemented *t*-tests (FDR threshold < 0.05) to compare the changes in expression and phosphorylation, as described previously^102^. Lists of upregulated or downregulated phosphoproteins (defined as having at least one significant upregulated or downregulated phosphopeptide) between two comparison groups were compiled and forwarded to ORA within KEGG^37^ and Reactome^70^ pathways using GSEApy^101^. There, the lists were compared to the background consisting of all quantified (phospho-)proteins. Pathways with an FDR threshold of 5% were considered significant (*p*-values were corrected using the BH^100^ method). To predict enhanced kinase activities The Kinase Library tool^71^ version 0.0.10 from PhosphoSitePlus^38^ was used and all sequence windows, together with corresponding FDR and Log_2_(FC) values were provided. These were compared to a background consisting of all quantified phosphopeptide sequence windows using one-sided Fisher’s exact tests. Values of entries that were exclusively explained by altered protein expression were replaced with default values to ensure that they did not distort the results (FDR = 1, Log_2_(FC) = 0). Predictions were required to have an adjusted *p*-value (BH^100^ corrected) < 0.05 and a dominant upregulated enrichment value ≥ 1.

### Histology

X-ray virtual histology was carried out using a commercial micro/nano-CT device, EasyTom XL Ultra (RxSolutions SAS, Chavanod, France). The scanner features a Hamamatsu reflection target microfocus X-ray source L10801. There exist two detectors, one flat panel detector with a thick, high-efficiency CsI converter, 127 µm pixel size, 1880 × 1494 pixels array, and 16-bit dynamic range. The other option is a CCD with 9 µm native pixel size and 4008 × 2672 pixels array. This CCD is used in 2 × 2 binning (18 µm) as the best physical resolution attainable is limited by its 20 µm thick Gadox scintillator. For the global scans the flat panel is used, and for the local scans presented here in **Fig. 5** the CCD is used. For **Fig. 5A** and **Fig. 5B** we used 1568 projections per scan, 90 kVp Tube acceleration voltage, and 110 µA tube current. For **Fig. 5C** we have SDD = 419.94 mm, SOD = 107.16 mm, Voxel size = 4.59 µm, 3.50 s exposure time, 6 frame averages per projection, and total scan time of 09:08 (hh:mm). In **Fig. 5A** we have SDD = 359.98 mm, SOD = 99.94 mm, Voxel size = 5.00 µm, 2.50 s exposure time, 6 frame averages per projection, 1568 projections per scan and scan time of 06:32 (hh:mm). The selected FFPE blocks were sliced every 2.5 µm using a Microm HM 355S (Theromo Scientific). Subsequently, the slices were stained by hematoxylin and eosin and scanned using a P250 scanner (3DHistech) to produce whole slide images in **Fig. 5B** and **Fig. 5D**.

## Data availability

Proteomics and Phosphoproteomics MS data are available within the PRIDE repository^103^ under the accession numbers: PXD055587 (Username, Password: cCTGfXZoX72j) and PXD055612 (Username, Password: gju2nnfDybrE).

## Acknowledgments

The authors gratefully acknowledge the FGCZ of the University of Zurich and ETH Zurich, and in particular Dr. Antje Dittmann and Laura Kunz, for the support on proteomics and phosphoproteomics analyses. **Fig. 1A** was precompiled with BioRender.com.

## Supporting information

Table S1: Sample description

Table S2: Differential expression analysis proteomics

Table S3: Differential expression analysis phosphoproteomics

Table S4: GSEA proteomics

Table S5: GSOA phosphoproteomics

Table S6: Enhanced kinase activities phosphoproteomics

**Figure S1.**
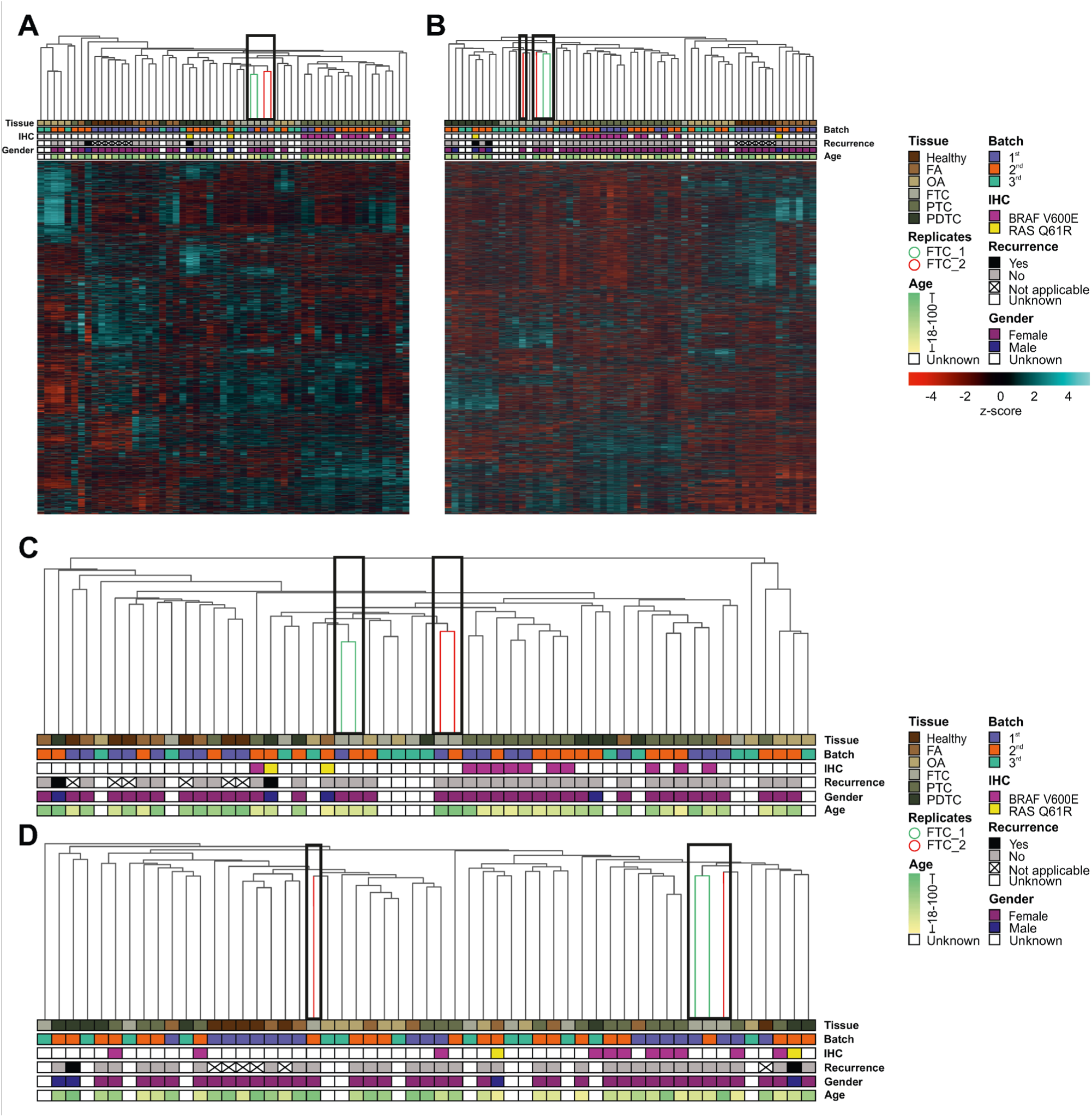
(related to Fig. 1): Hierarchical clustering. **A, B** Hierarchical clustering (HC) with information on tumor subtype, batch, immunohistochemistry (IHC) staining, recurrence, gender, and age of (**A**) 2’540 proteins and (**B**) 5’618 phosphopeptides (extended version of **Fig. 1B-C**). Rows and columns are clustered, while only the column dendogram is displayed. Row intensities are z-score scaled. Biological replicates of two FTC samples (FTC_1 marked green and FTC_2 red in the dendogram) are highlighted with black boxes. **C, D** HC of (**C**) 2’540 proteins and (**D**) 5’618 phosphopeptides with unbiased imputation, where missing data were sample-wise randomly sampled from a constructed normal distribution with a negative shift from the mean of the actual measured entries without taking the experimental design (subtype-identity) into consideration. Intensities of individual entries are not displayed. Biological replicates of two FTC samples (FTC_1 marked green and FTC_2 red in the dendogram) are highlighted with black boxes.

**Figure S2.**
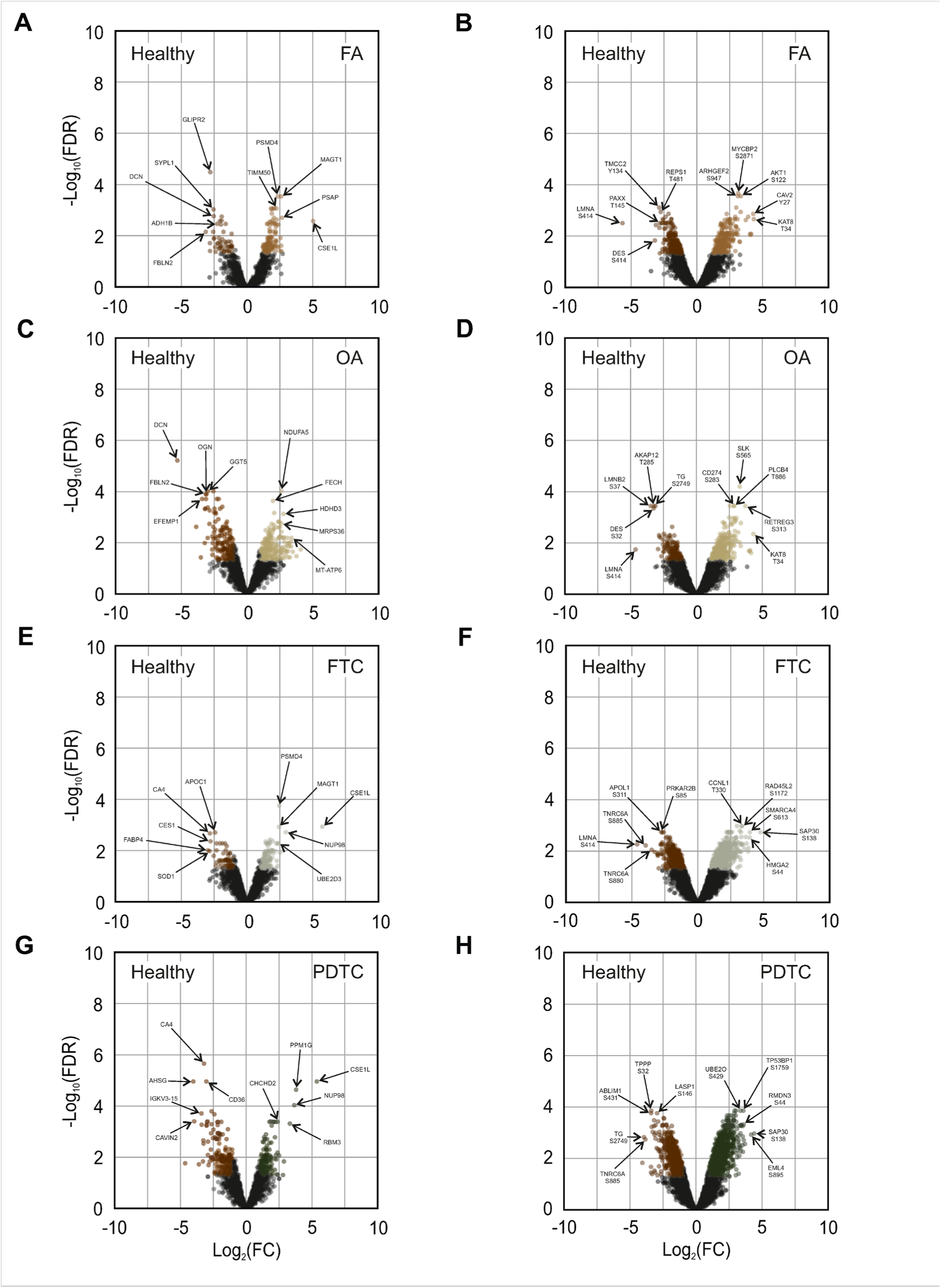
(related to Fig. 2): Volcano plots comparing protein expression and phosphorylation between thyroid neoplasms and healthy control tissue. **A, B** Volcano plots of quantified (**A**) proteins and (**B**) phosphopeptides comparing FA with healthy thyroid control tissues. **C, D** Volcano plots of quantified (**C**) proteins and (**D**) phosphopeptides comparing OA with healthy thyroid control tissues. **E, F** Volcano plots of quantified (**E**) proteins and (**F**) phosphopeptides comparing FTC with healthy thyroid control tissues. **G, H** Volcano plots of quantified (**G**) proteins and (**H**) phosphopeptides comparing PDTC with healthy thyroid control tissues. Colored entries represent significant hits with FDR < 0.05 and absolute Log_2_(FC) ≥ 1. Top five up– and downregulated hits were annotated based on ranking according to –Log_10_FDR × Log_2_(FC).

